# The Endothelin Receptor Antagonist Macitentan Ameliorates Endothelin-Mediated Vasoconstriction and Promotes Neuroprotection of Retinal Ganglion Cells in Rats

**DOI:** 10.1101/2020.10.27.357814

**Authors:** Dorota L. Stankowska, Wei Zhang, Shaoqing He, Vignesh R. Krishnamoorthy, Payton Harris, Trent Hall, Renuka M. Chaphalkar, Bindu Kodati, Sai H. Chavala, Raghu R. Krishnamoorthy

## Abstract

**Purpose:** To determine if dietary administration of the dual ET_A_/ ET_B_ receptor antagonist, macitentan, could protect retinal ganglion cells (RGCs) following endothelin-1 (ET-1) mediated vasoconstriction in Brown Norway rats.

**Methods:** Adult male and female Brown Norway rats were either untreated or treated with macitentan (5 mg/kg body weight) once a day for 3 days followed by intravitreal injection of either 4 μl of 500 μM ET-1 (2 nmole/eye) or vehicle in one eye. Imaging of the retinal vasculature using fluorescein angiography was carried out at various time points, including, 5, 10, 15, 25 and 30 minutes. Following the imaging of the vasculature, rats were either treated with macitentan (5 mg/kg/body weight in dietary gels) or untreated (control gels without medication). Following treatments, rats were euthanized, retinal flat mounts were prepared, immunostained for RGC marker Brn3a, imaged and surviving RGCs were counted in a masked manner.

**Results:** Vasoconstrictive effects following intravitreal ET-1 injection were greatly reduced in rats administered with macitentan in the diet prior to the ET-1 administration. ET-1 intravitreal injection produced a 31% loss of RGCs which was significantly reduced in macitentan-treated rats. Following ET-1 administration, GFAP immunostaining was increased in the ganglion cell layer as well as in the retrolaminar region, suggestive of astrocytic activation by ET-1 administration. RGC numbers in macitentan treated and ET-1 injected rats were similar to that observed in control retinas.

**Conclusions:** ET-1-mediated neurodegeneration could occur through both vascular and cellular mechanisms. The endothelin receptor antagonist, macitentan, has neuroprotective effects in retinas of Brown Norway rats that occurs through different mechanisms, including, enhancement of RGC survival and reduction ET-1 mediated vasoconstriction.

## Introduction

Endothelins are a family of potent 21-amino-acid vasoactive peptides which play an important role in the maintenance of vascular tone in the cardiovascular system [1, 2] **Figure 1A**. There are three isoforms of endothelins encoded by separate genes: endothelin-1 (ET-1), endothelin-2 (ET-2) and endothelin-3 (ET-3) [3]. Several studies have shown that endothelins contribute to the pathophysiology of glaucoma [4]. ET-1 levels are elevated in the aqueous humor of primary open angle glaucoma (POAG) patients [5, 6] and also in animal models of glaucoma [7–9]. In particular, endothelin B (ET_B_) receptors have been shown to play a key role in neurodegeneration in animal models of glaucoma [10–12], **Figure 1B**. More recently, our laboratory demonstrated the upregulation of the ET_A_ receptor in a rat model of ocular hypertension [13]. While the role of the ET_A_ receptor in neurodegeneration is not completely understood, studies have shown that the inhibition of both endothelin receptors (ET_A_ and ET_B_) provides neuroprotection in an inheritable mouse model of glaucoma [14]. ET⍰1 promotes vasoconstriction by interacting with endothelin receptors ET_A_ and ET_B_, hence the potential mechanisms responsible for ET-1 actions could involve a vascular component, mostly as decreased blood flow to the optic nerve head. The vascular dysregulation, besides the intraocular pressure (IOP) elevation is also thought to contribute to the glaucoma pathophysiology [15–17]. The current study was aimed at determining if a dual endothelin antagonist macitentan could attenuate retinal vasoconstriction, promote RGC protection following an intravitreal administration of an acute dose of ET-1 (4μl of 500 μM, 2 nmole/eye).

**Figure. 1.**
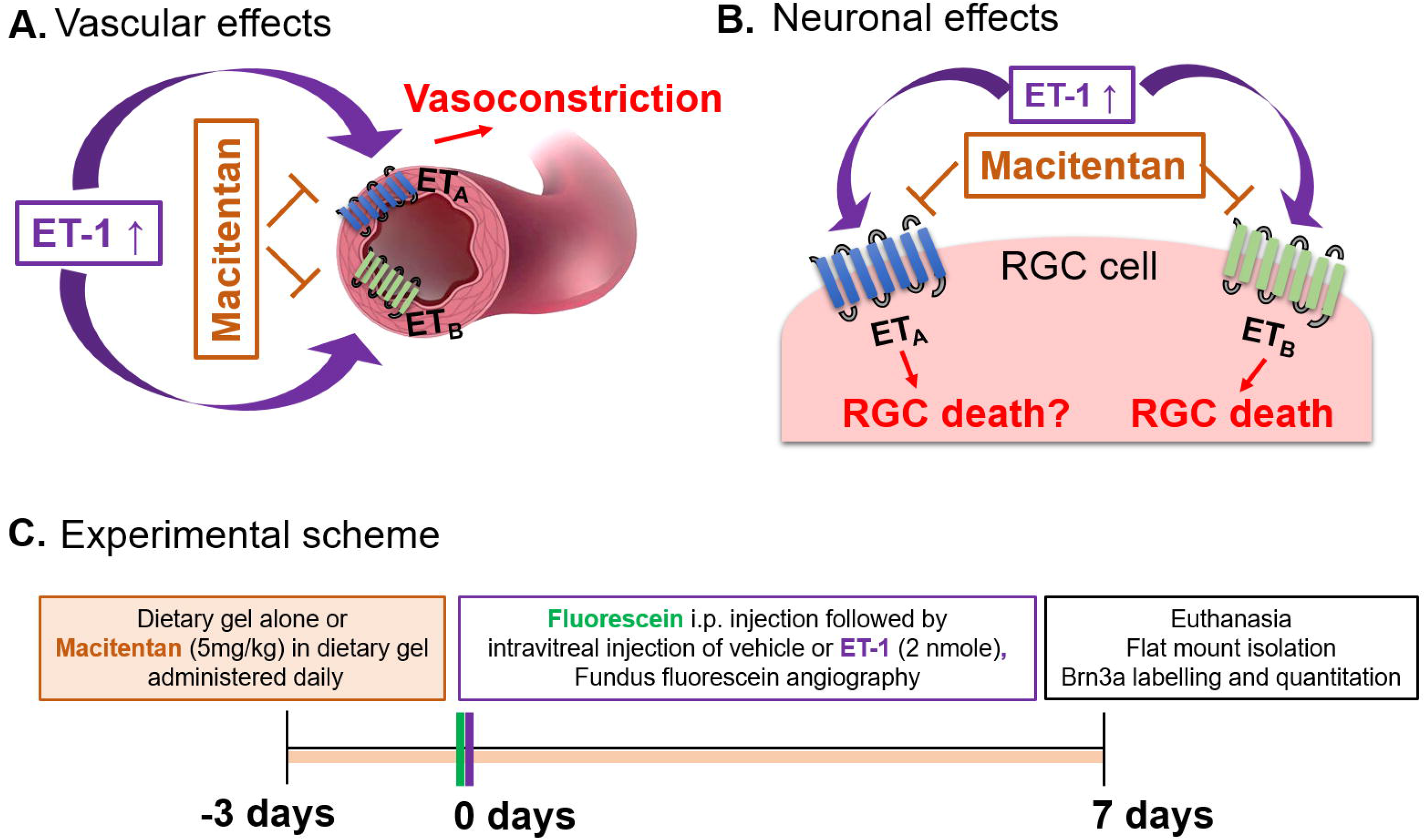
Hypothetical mechanisms of action of endothelin 1 (ET-1). **(A)**Vascular effects of ET-1, **(B)** Effects of ET-1 on the retinal ganglion cells. **(C)**Experimental scheme.

## Methods

### Animals

Animal studies were performed in accordance with the Association for Research in Vision and Ophthalmology (ARVO) resolution for the Use of Animals in Ophthalmic and Vision Research and approved by the University of North Texas Health Science Center (UNTHSC) Institutional Animal Care and Use Committee (IACUC). Retired breeder male and female Brown Norway rats (8- to 11-months-old) were obtained from Charles River (Wilmington, MA).

### Macitentan

Macitentan was a kind gift from Actelion Pharmaceuticals US, Inc. (CA, USA). Macitentan treatment (5 mg/kg body weight/day) was initiated three days prior to the ET-1 or vehicle intravitreal injections and was continued for another 7 days post-intravitreal injection (for a total 10 days) (**Figure 1C)**. To ensure even consumption, macitentan was administered orally by mixing the drug into DietGel^®^ Recovery (Clear H_2_0, Westbrook, ME). Untreated Rats were fed with the DietGel^®^ alone. Rats were monitored to ensure complete consumption of the medication.

### Intravitreal Injections of ET-1 or Vehicle

Endothelin-1 (Bachem, Torrance, CA, USA) was resuspended in a vehicle solution (0.25% acetic acid adjusted to pH 7.0 with sodium hydroxide) to final concentration of 500 μM. Intravitreal injections were performed using a Hamilton syringe with a 32-gauge needle. The rats were first anesthetized by intraperitoneally injecting an anesthetic cocktail of ketamine (55 mg/kg) /xylazine (5.5 mg/kg) /acepromazine (1.1 mg/kg). A single drop of 0.5% proparacaine hydrochloride (Alcon Laboratories, Inc., Fort Worth, TX, USA) and 1% tropicamide were applied to both eyes. Intravitreal injection of 4μl of 500 μM ET-1 or vehicle was carried out in one eye of Brown Norway rats (with continuous observation of the needle in the center of the vitreous cavity to avoid lens injury). The injections were performed through the sclera, approximately 1 mm behind the limbus and solutions were slowly (~30 s) injected into the vitreous chamber of the eye. To prevent the injected solution from escaping the eye, the needle tip was held in the eye for 30 seconds and then gradually withdrawn. Triple antibiotic was applied to the site of injection to prevent infections and allowing healing to occur.

### Fundus photography (FP) and fundus fluorescein angiography (FFA)

Rats were anesthetized with a cocktail of ketamine (55 mg/kg)/xylazine (5.5 mg/kg)/acepromazine (1.1 mg/kg) and pupils were dilated using eye drops of a mixture of 0.5% tropicamide and 0.5% phenylephrine hydrochloride. The cornea surface was anesthetized by 0.5% proparacaine hydrochloride and was kept moist with GenTeal Tears lubricant eye gel (Alcon Laboratories Inc., Fort Worth, TX). AK-FLUO (10%, Akorn Inc., Lake Forest, IL) was injected intraperitoneally at 1.5 μl/g of body weight. ET-1 was intravitreally injected 3 minutes following administration of fluorescein and imaging of the retina was carried out. Micron IV retinal imaging microscope (Phoenix Research Laboratories, Pleasanton, California) was used to capture the fundus images and fluorescein angiography images. Serial images were then captured at different time points including, 5, 10, 15, 20, 25 and 30 minutes following ET-1 injection. The body temperature was maintained at 37°C using a heating pad. Following recovery from anesthesia, animals were provided food and water ad libitum.

### Retinal Flat Mount Immunostaining

Following ET-1 injections, rats were fed with either macitentan (5 mg/kg body wt) in dietary gels or dietary gels alone (untreated control) for seven days. Following the treatments, animals were euthanized, eyes were enucleated. The eye cups were fixed overnight at 4°C in 4% paraformaldehyde (PFA) and retinal flat mounts were prepared. To prevent non-specific binding with the secondary antibody, blocking was carried out using 5% normal donkey serum in 5% BSA overnight at 4°C. Retinal flat mounts were incubated in primary antibody solution, goat anti-Brn3a (1:200; Santa Cruz) for 72 hours at 4°C. Subsequently, secondary antibody incubation was carried out using a 1:1000 dilution of Alexa 647 conjugated donkey anti-goat antibody (Life Technologies, Carlsbad, CA) overnight at 4°C. The retinal flat mounts were mounted on slides using Prolong Gold anti-fade (Life Technologies). All images were taken in the Cytation-5 microscope (BioTek, Winooski, VT).

### Semi-Automatic Retinal Ganglion Cell Counting

The images of immunostained retinal flat mounts were uploaded to ImageJ, a free photo editor designed for biology research (Rasband, 1997-2018). The images were then converted to 8-bit greyscale to reduce background noise and processed with the automatic nuclei counter plugin “ICTN” which automatically counts high contrast points within the image. To maintain consistency in cell counts the ICTN settings were set to detect cells of a specific width, distance apart, and contrast threshold (8, 6, and 1.5, respectively). The cells not detected by the ICTN program were then counted manually by a masked observer and summed as total RGC counts.

### Primary optic nerve head astrocytes

Rat primary optic nerve head astrocytes were isolated from adult 8 to 10 week old female Sprague-Dawley rats and cultured using AM-a medium (ScienCell Research Laboratories, Inc., CA, USA, Catalog #1801) with 2% FBS [18]. When cells reached the confluence at 80%, the culture medium was changed to serum-free astrocyte medium. Pretreatment of the cells were carried out by treatment with 1μM macitentan or 0.1% DMSO (vehicle) for 30 min. After the pretreatment, the cells were treated with ET-1 at a final concentration 100 nM for 24 hours.

At the end of the treatment, the cells were harvested using TRIZOL and the total RNA was extracted following the manufacturer’s instruction. The quantity and quality of total RNA was monitored by Nanodrop 2000 (Fisher Scientific). cDNA was synthesized using BioRad iScript Reverse Transcription Supermix (#1708840), and qPCR was performed to detect the expression of GFAP and fibronectin using BioRad SsoAdvance Universal SYBR Green Supermix (#1725271). Detection of cyclophilin served as an internal control. The following primers were used for the qPCR analyses:

Rat GFAP (Sense): 5’-AAATTGCTGGAGGGCGAAGA
Rat GFAP (Anti-sense): 5’-CCGCATCTCCACCGTCTTTA
Rat Fibronectin (Sense): 5’-AAACCGGGAAGAGCAAGAGG
Rat Fibronectin (Anti-sense): 5’-CCTAGGTAGGTCCGTTCCCA
Rat Cyclophilin A (Sense): 5’-CGGAGAGAAATTTGAGGATGA
Rat Cyclophilin A (Anti-sense): 5’-CATCCAGCCACTCAGTCTTG

### Statistical analysis

SigmaPlot software (Systat Software, Inc., San Jose, CA, USA) was used for statistical analyses. The data are expressed as the mean ± standard error (SEM) of three or more independent experiments. Statistical significance of experiments was determined by using ANOVA on ranks with the Dunn’s Method test or one way ANOVA with Bonferroni t-test at p ≤ 0.05.

## Results

### Vasoconstrictive effects of intravitreally injected ET-1 are reduced in rats administered with macitentan in the diet

To test the ability of the dual ET_A_/ET_B_ antagonist, macitentan on ET-1 mediated vascular changes in the eye, we treated male and female Brown Norway rats with either the dietary gel alone or dietary gels containing macitentan (5mg/kg), starting 3 days prior the ET-1 intravitreal injection. On the day of the experiment (day “0”) the rats were intraperitoneally injected with sodium fluorescein to visualize retinal vasculature. Rats were intravitreally injected either with the 4 μl of vehicle (n=6) or with 4μl of 500μM ET-1 (n=6). Images were captured at various time points including: 0, 5, 10, 15, 20, 25 and 30 min following the intravitreal injections. We observed profound vasoconstriction (particularly in the retinal arteries) in ET-1 injected eyes (**Figure 2A**), which was reflected by a significant decrease in vessel diameter and vascular density particularly at 10 minutes following ET-1 administration (**Figure 2B-E**). Rats treated with the ET_A_/ET_B_ antagonist macitentan exhibited significantly lesser vasoconstriction, which was reflected in higher vessel diameter and vascular density, compared to the untreated rats injected with ET-1. The significant constriction was evident as early as 5 min following ET-1 injection and lasted up to 30 minutes. On the other hand, vasoconstriction was delayed (particularly evident at the site of injection) and much lesser vasoconstriction was observed at 10 min in rats administered macitentan in dietary gels, compared to that in untreated rats (**Figure 2B-E**). These findings suggest that blocking both of the endothelin receptors partially prevents vasoconstriction or appreciably delays its onset. Rats injected with ET-1 and fed with dietary gels alone, showed no sign of recovery from vasoconstriction up to 30 min (Supplementary Fig. S1). There were no changes in vasoconstriction in rats injected with vehicle at any of the tested time-points (for both groups: dietary gel or macitentan administered).

**Figure. 2.**
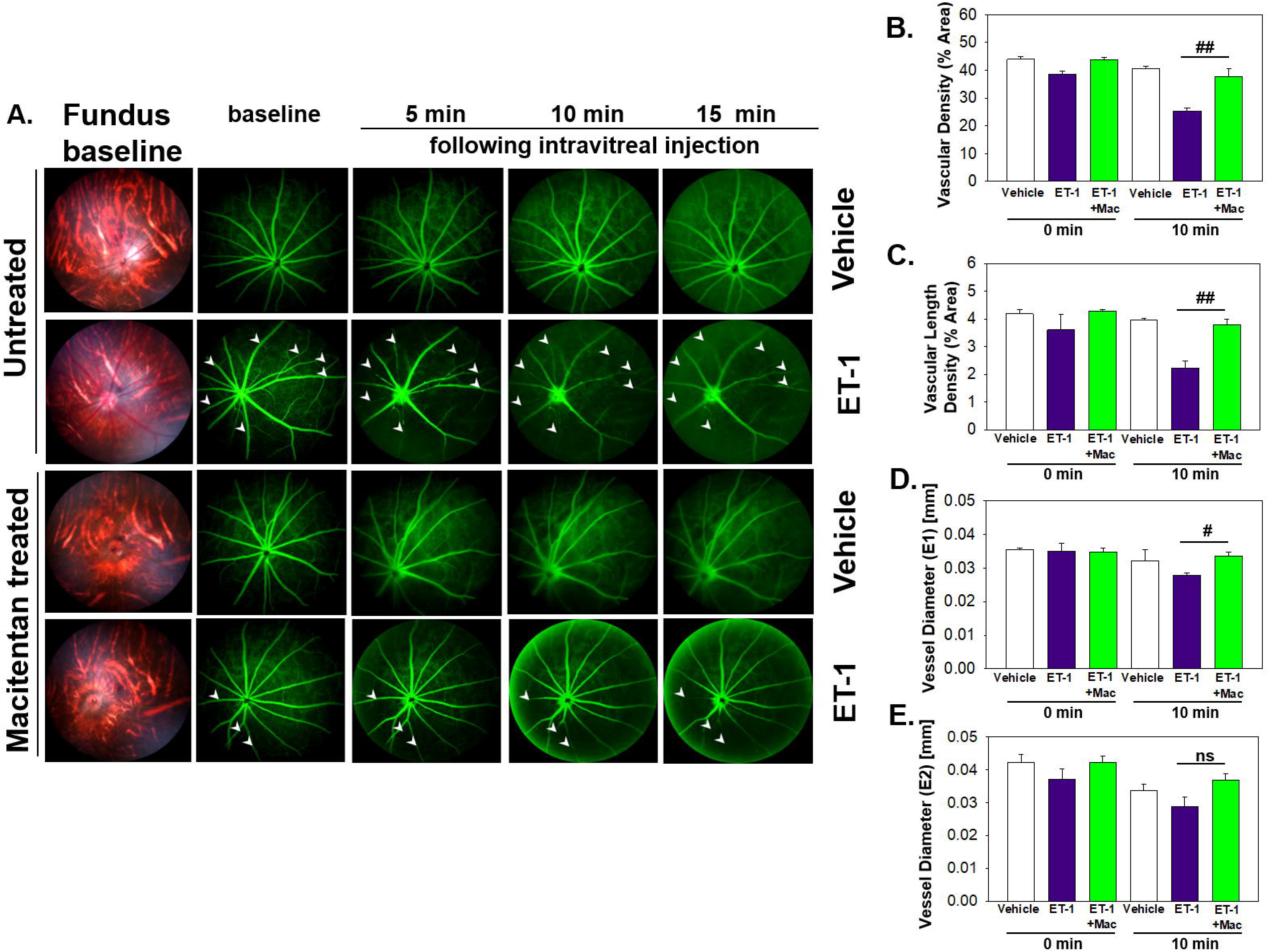
Retinal vasculature fluorescein angiography and analysis of vasculature changes in Brown Norway rats. Rats were either treated or untreated with macitentan for three days prior to intravitreal injection of ET-1. **(A)**The retinal vasculature was imaged with the Micron IV microscope prior to ET-1 injection (baseline) and subsequently at 5, 10 and 15 minutes. Arrowheads indicate the blood vessels which had the most prominent vasoconstriction. **(B)**The vascular density was measured using FIJI Vessel Analysis plugin and presented as vessel area/total area * 100%, n= 3 rats per treatment. **(C)**The vascular density length was assessed with FIJI Vessel Analysis plugin and graph represents skeletonized vessel area/total area * 100%, n= 3 rats per treatment group. **(D, E)**The vessel diameter was calculated using FIJI Vessel Analysis plugin at two distances form the optic nerve head at eccentricity 1 (E1, **D**) and eccentricity 2 (E2, **E**), at least 3 rats per group were analyzed. Data on the graphs is presented as Mean ± SEM, where (##) = p<0.05 using all pairwise multiple comparison procedures (Dunn’s Method), (#) = p< 0.05 using all pairwise multiple comparison procedures (Bonferroni t-test) (ns) = no statistical difference.

### ET-1 intravitreal injection produced RGC loss which was significantly attenuated in macitentan-treated rats

To determine if ET-1 treatment promoted RGC loss and the effect of macitentan on RGC survival, macitentan treatment was continued one week following ET-1 or vehicle injection to one eye of the rats. Following the treatments, rats were euthanized and retinal flat mounts were prepared as described in **Figure 1C**. Labelling of retinal ganglion cells was performed on retinal flat mounts as described previously with minor modifications using RGC marker Brn3a, which label surviving RGC [19] (**Figure 3A**). Semi-automatic counts were performed in all experimental groups including a naïve group in which no injection or treatment was carried out. As seen in **Figure 3A** naïve RGCs have a good morphology and are brightly stained using Brn3a antibody. In contrast, retinas treated with ET-1 show tremendous decrease in Brn3a immunostaining which was attenuated in ET-1 and macitentan treated rats. Vehicle injected eyes show partial RGC loss possibly due to mild inflammation following the invasive procedure of injection. **Figure 3B** illustrates average number of RGCs per each field of view. As expected naïve rats have very high RGC numbers (4693±90). ET-1 injection significantly reduced that number by 31% (2585±271) and macitentan reduced the cell loss to the level of naïve (control) animals (4736±239). Vehicle rats showed a decline in RGC number (3255±243).

**Figure. 3.**
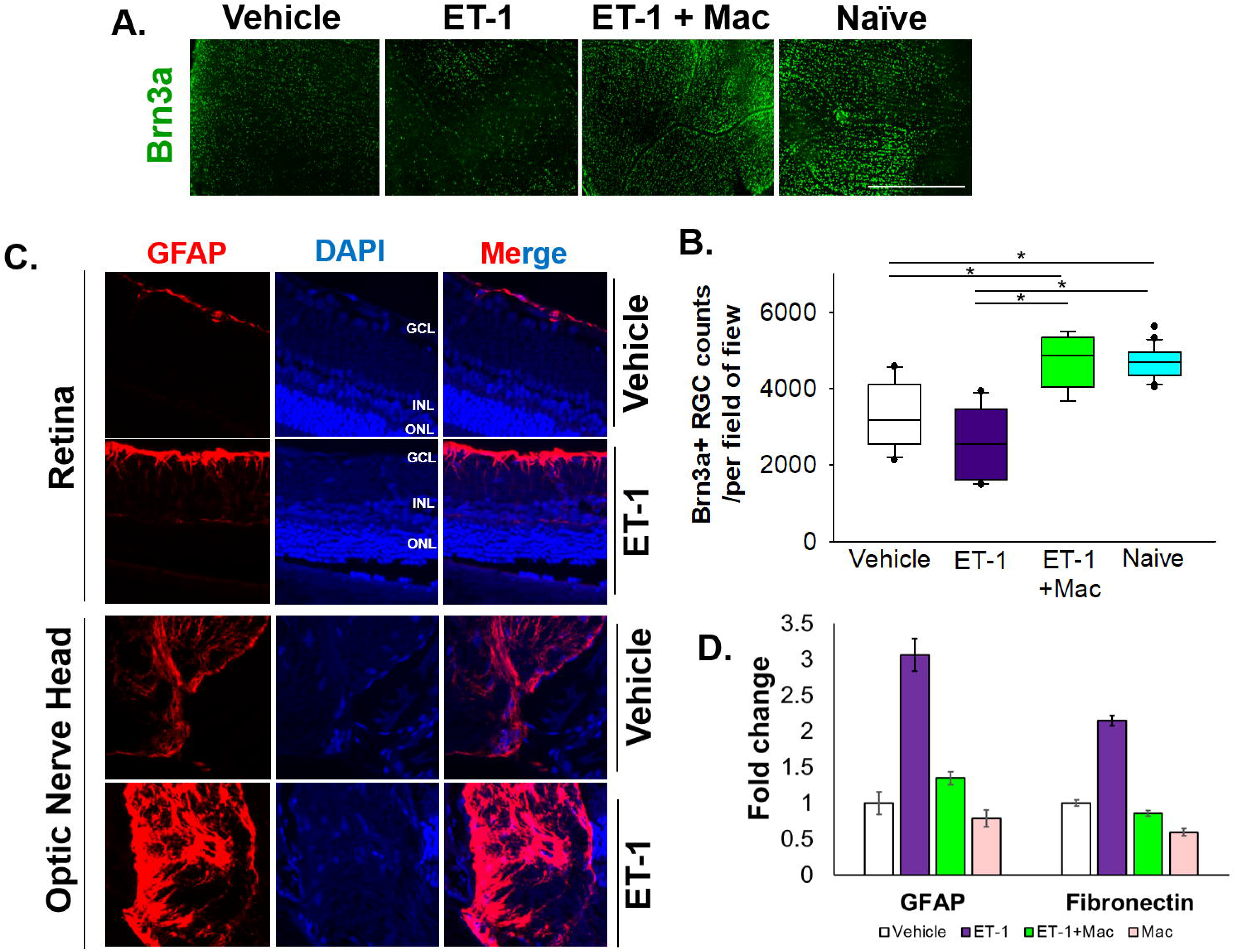
Treatment with macitentan significantly reduces ET-1 mediated RGC loss in Brown Norway rats and ameliorates activation of optic nerve astrocytes. **(A)**Brown Norway rats were untreated or treated with macitentan (Mac) for 3 days following which they were intravitreally injected with ET-1 or vehicle. Macitentan (or control gel) treatments were continued for additional 7 days. Rats were then sacrificed and retinal flat mounts were isolated. The panel shows representative images of retinal flat mounts immunostained with an antibody to the RGC marker Brn3a. Scale bar represents 1000 μm **(B)**A plot illustrating average number of Brn3a-positive RGCs per field of view (4-8 images per each retina were acquired, n=3-5 rats per group, n=12 for naïve group). Bars represent mean ± SEM. (*) = p<0.05 using all pairwise multiple comparison procedures (Dunn’s Method). **(C)**Immunohistochemical analysis of GFAP expression in retinas and optic nerves of Brown Norway rats 24 hours following intravitreal injection of either vehicle or ET-1. GCL - ganglion cell layer, INL – inner nuclear layer, ONL – outer nuclear layer, DAPI indicated cell nuclei. **(D)**Quantitative PCR analysis of expression of GFAP and Fibronectin in rat primary optic nerve head astrocytes isolated from adult Sprague-Dawley rats following 24 hour treatment with either the vehicle, ET-1, Macitentan or a combination of ET-1 and macitentan. The graph represents the mean ± SD of one representative experiment. The same experiment was repeated 3 times and similar trend was observed.

### ET-1 treatment produced an upregulation of GFAP in rat retinas as well as primary cultures of optic nerve head astrocytes

Astrocytes play a pivotal role in maintaining the cellular environment of retinal neurons. To determine if ET-1 could produce activation of astrocytes (since they are in the vicinity of RGCs), we tested retina sections from Brown Norway rats, 24 hours post-intravitreal injection of 2 nmole of ET-1. As seen in **Figure 3C**, an increased immunostaining for GFAP was observed in the nerve fiber layer as well as in the optic nerve head of the rats injected with ET-1, compared to vehicle-injected rats.

Similar to the *in vivo* experiments, cultured primary optic nerve head astrocytes treated with ET-1 (100 nM, 24 hours) showed a similar 3-fold increase in mRNA levels of GFAP, as determined by a q-PCR analysis. Treatment with the ET_A_/ET_B_ dual antagonist, macitentan, completely blocked ET-1 mediated increase in GFAP expression. In addition, we also observed a 2-fold increase in mRNA expression of fibronectin, which was prevented by treatment with macitentan. Taken together, ET-1 acting through its receptors has the ability to promote activation of astrocytes resulting in a pro-fibrotic phenotype, which could have damaging effects on RGCs and their axons.

## Discussion

Endothelins (including ET-1, ET-2 and ET-3) have been shown to be key players promoting neurodegeneration in glaucoma [20–23]. Endothelins could act through both vascular and cellular mechanisms to promote degeneration in glaucoma. These could occur by hypoxic mechanisms and/or activation of a pro-apoptotic signaling thereby, compromising the survival of RGCs and their axons. Vascular mechanisms are difficult to assess in glaucoma patients due to the inter-individual variations in vascular architecture and perfusion of the optic nerve head. Polak et al., (2003) found a decrease in blood flow both in the optic nerve head as well as choroidal in healthy human subjects following intravenous administration of ET-1, which were blocked by co-administration of an ET_A_ receptor antagonist [38]. Several previous studies have shown that continuous perfusion of ET-1 at the optic nerve head could produce a decline in optic nerve head blood in rats, rabbits and monkeys [24–26]. However, most of these studies did not assess both vascular changes as well as cellular loss of RGCs and additionally did not test the effect administration of an endothelin receptor antagonist.

Intravitreal administration of ET-1 into rat eyes can promote loss of RGCs acting through endothelin receptors [27, 28]. These studies using ET_B_-receptor deficient rats, demonstrated the causative role of the ET_B_ receptor in loss of RGCs, however, the ability of a endothelin receptor antagonist to protect RGCs was not demonstrated in rats. Kiel (2000) found that administration of ET-1 in rabbits decreased choroidal blood flow which was blocked by the non-selective antagonist A-182086, while the ET_A_ antagonist (FR-139317) enhanced the dilation and blocked the constriction [29]. These findings raised the possibility of using endothelin receptor antagonists as a neuroprotective agent in glaucoma.

ET-1 has a relatively short half-life (<5 min) in circulation [30]. Hence, the plasma levels of ET-1 may not be reflective of its actual peak concentrations both in the circulation as well as in the aqueous humor. Nevertheless, many publications have implicated an increase in ET-1 concentrations in the aqueous humor as well as plasma of glaucoma patients, compared to age-matched control subjects. Most studies in culture are unable to detect ET-1 mediated cellular effects at concentrations below 10 to 100 nM ET-1.

Chauhan and colleagues perfused ET-1 through osmotic mini-pumps at the retrobulbar region of the rat eye, for various time durations from 21 to 84 days and found a time-dependent loss of RGCs and damage to their axons [26]. Similar to our findings, Lau and colleagues demonstrated [31] 25 to 44% RGC loss (at 1 and 4 weeks respectively) was observed after intravitreal administration of ET-1. This suggests that irrespective of the site of administration, ET-1 could produce neurodegeneration of RGCs and their axons. The precise mechanisms underlying ET-1 mediated neurodegeneration of RGCs are not completely understood. Studies from cardiovascular research have provided some insight into mechanisms by which activation of endothelin receptors could produce cellular damage, some of which may be relevant to neurodegeneration. For instance, perfusion of sarafotoxin (an ET_B_ receptor agonist) was found to greatly elevated superoxide production in the sensory ganglia as well as in glial cells. ET-1 acting through the ET_B_ receptor produced efflux of glutamate from cultured brain astrocytes, suggestive of its ability to promote excitotoxic effects [32]. Previous work from our lab demonstrated that IOP-mediated RGC loss and optic nerve degeneration were significantly attenuated in ET_B_ receptor-deficient rats [12]. McGrady showed that ET_A_ receptors are also elevated in the retinas of rats with elevated IOP and have the ability to upregulate ET_B_ expression [13]. We are now showing that ET-1 intravitreal injection produces activation of rat optic nerve head astrocytes in vivo (indicated by GFAP staining). Taken together, blocking both ET_A_ and ET_B_ receptor would be beneficial to halt ET-1 mediated neurodegeneration occurring through both vascular and cellular mechanisms. In fact, Howell and colleagues demonstrated that bosentan (dual antagonist of ET_A_ and ET_B_ receptor) treatment promoted neuroprotection of optic nerve axons in the congenital DBA/2J model of glaucoma in mice [33]. However, the authors did not assess the effects of bosentan on vascular changes in the retina following endothelin administration. Macitentan has higher potency and efficacy (due to higher receptor occupancy times) as an endothelin receptor antagonist, compared to bosentan [34, 35]. In a subsequent study, Howell et al., (2014) found appreciable axoprotection in 80% of 10 month old DBA/2J mice treated with macitentan (30 mg/kg body wt). In addition, the authors used bosentan (100 mg/kg body wt) in C1q knockout DBA/2J mice to show appreciable neuroprotection of optic nerve axons in 80% of 12 month old animals [36]. However, the authors did not report any enhancement in RGC survival, which is reported in this study.

In summary, the novel aspects of this study include the use of a lower dose of macitentan (5 mg/kg) to generate protective effects including, amelioration of retinal vasoconstriction, enhanced RGC survival, as well as reduction of optic nerve head astrocyte activation. Studies have shown that use of a similar dose in humans did not produce any major adverse effects, minor side-effects including headache and back pain during this treatment were also observed in the placebo group [37]. We are also demonstrating that treatment of rat optic nerve head astrocytes with macitentan produces a decreases in ET-1 mediated mRNA expression of some markers of astrocytes activation, namely GFAP and fibronectin. Findings from the current study have major implications for the use of orally administered macitentan (which is FDA approved for use in pulmonary hypertension) as a neuroprotective adjunct therapy to IOP lowering drugs, for the treatment of glaucoma. Additionally, macitentan has the potential to combat vasoconstrictive effects observed in acute angle closure glaucoma, central retinal artery/vein occlusion and related ischemic disorders of the retina.

## Conclusions

1. Vasoconstrictive effects following intravitreal ET-1 injection were greatly reduced in rats administered with macitentan in the diet prior to the ET-1 administration.
2. ET-1 intravitreal injection produced a 31% loss of RGCs which was significantly lower in macitentan-treated rats. RGC counts following ET-1 injection and macitentan treatment were similar to that observed in control retinas.
3. Macitentan has neuroprotective effects in retinas of Brown Norway rats that possibly occur through different mechanisms, including, enhancement of RGC survival, decreasing activation of optic nerve head astrocytes and the reduction of ET-1 mediated vasoconstriction and preventing ischemia.

## List of abbreviations

ET-1: Endothelin-1
RGC: Retinal Ganglion Cell
ET_A_: Endothelin A Receptor
ET_B_: Endothelin B Receptor

## Declarations

### Ethics approval

All experiments and procedures performed in this study were in agreement with the ARVO guideline for the use of animals in research and were approved by the Institutional Animal Care and Use Committee (IACUC) at the UNT Health Science Center, Fort Worth.

### Consent for publication

All authors have reviewed the manuscript and given their consent for publication.

### Availability of data and material

Not applicable. Macitentan was a kind gift from Acetelion Pharmaceuticals which was obtained through a Material Transfer Agreement (MTA).

---

**Figure. S1.**
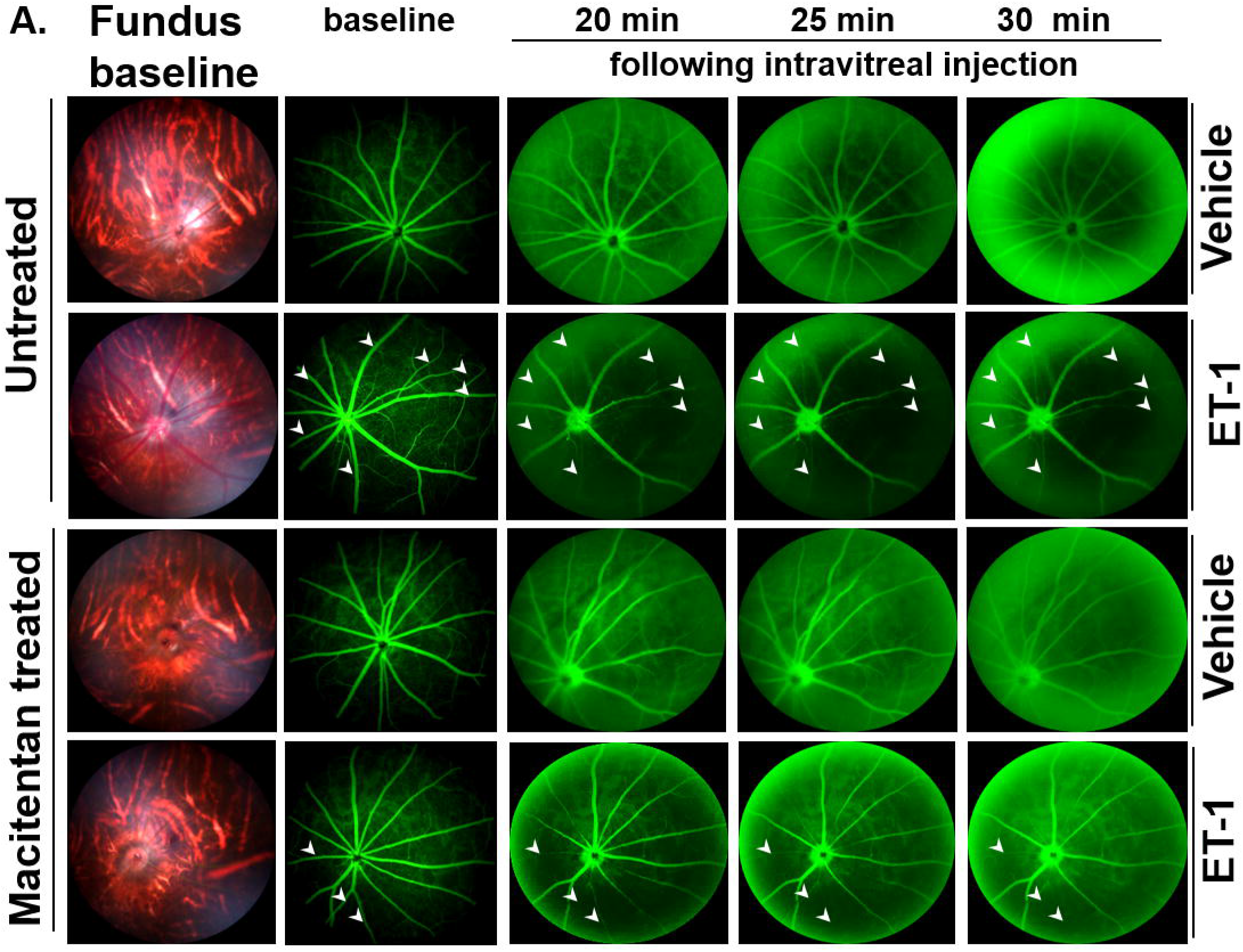
Vasculature fluorescein angiography of the retina in Brown Norway rats. Rats were either treated or untreated with macitentan for three days prior to intravitreal injection of ET-1. The retinal vasculature was imaged with the Micron IV microscope directly before the ET-1 injection (indicated as baseline) and continued to capture at 20, 25 and 30 minutes, n=3-5 rats per group. Arrowheads indicate the blood vessels which had the most noticeable vasoconstriction.

